# Living Neural Networks: Dynamic Network Analysis of Developing Neural Progenitor Cells

**DOI:** 10.1101/055533

**Authors:** Arun S. Mahadevan, Nicolas E. Grandel, Jacob T. Robinson, Kevin R. Francis, Amina A. Qutub

## Abstract

The architecture of the mammalian brain has been characterized through decades of innovation in the field of network neuroscience. However, the assembly of the brain from progenitor cells is an immensely complex process, and a quantitative understanding of how neural progenitor cells (NPCs) form neural networks has proven elusive. Here, we introduce a method that integrates graph-theory with long-term imaging of differentiating human NPCs to characterize the evolution of spatial and functional network features in NPCs during the formation of neural networks in vitro. We find that the rise and fall in spatial network efficiency is a characteristic feature of the transition from immature NPC networks to mature neural networks. Furthermore, networks at intermediate stages of differentiation that display high spatial network efficiency also show high levels of network-wide spontaneous electrical activity. These results support the view that network-wide signaling in immature progenitor cells gives way to a hierarchical form of communication in mature neural networks. We also leverage graph theory to study the spatial features of individual cell types in developing cultures, uncovering spatial features of polarized neuroepithelium. Finally, we employ our method to uncover aberrant network features in a neurodevelopmental disorder using induced pluripotent stem cell (iPSC) models. The “Living Neural Networks” method bridges the gap between developmental neurobiology and network neuroscience, and offers insight into the relationship between developing and mature neural networks.

## INTRODUCTION

The study of complex, multiscale brain networks using concepts from graph theory and network science – an approach collectively termed network neuroscience – has enabled significant insight into the structural and functional organization of the brain^1,2^. Micro-connectomics – the study of organizational principles of neuronal networks at the cellular scale^3,4^, is an important subset of complex brain networks that has yielded insight into architectural features of the nervous system at the level of their basic building blocks. A number of studies have applied graph-theoretic approaches to study the functional and anatomical connectivity of *in vitro* neuronal cultures formed from dissociated cells^5–7^. These studies have shown, for instance, that *in vitro* neuronal networks self-assemble in a small-world topology with high clustering and low path length. Micro-connectomic analyses of slice cultures^8,9^, and more recently, large-scale network reconstructions^10^, have revealed fundamental properties of mammalian cortical networks, such as long-tailed synaptic connectivity, the presence of preferentially connected subgroups of neurons and overrepresentation of certain network motifs. However, despite the progress made in understanding the fundamental architectural features of neural networks, no models have been available to study the development of human neural networks from progenitor cells in a quantitative manner, nor have there been tools to characterize the spatial and functional dynamics of network formation at the cellular level. Cell-cell communication among neural progenitor cells (NPCs) is an essential aspect of human nervous system development. Neural progenitor cells cluster together in specialized microenvironments or niches where communication with neighboring cells plays an important role in determining cell behavior^11^. Prior to the formation of functional synapses, NPCs display structured intercellular communication that plays a critical role in the spatiotemporal control of self-renewal and differentiation, and shapes developing neural circuits. Examples of structured cell-cell communication include patterned, spontaneous electrical activity mediated partly through gap junctional coupling^12–14^, maintenance of intercellular configurations through tight junction proteins^15^ and control of cell differentiation through Notch signaling^16,17^. Notably, the predominant forms of communication employed by NPCs can be described as juxtacrine signaling, i.e., requiring direct cell-cell contact. This is in contrast to communication in mature neuronal networks, where the physical wiring among neurons is important and network-wide information is conveyed primarily through synaptic contacts. Given the dominance of juxtacrine modes of signaling among progenitor cells, graph-theoretic approaches need to be modified to study developing neural networks.

To enable the bridge between development and network neuroscience, we harness another emerging technology: directed differentiation of human stem cells. In recent years, significant advances in stem cell differentiation protocols have made it possible to produce a multitude of neuronal types and have provided a standardized workflow for generating functional human neurons *in vitro*^18^. Monolayer cultures of human stem cells capture many aspects of *in vivo* neural development, such as spatial and temporal features of cortical neurogenesis^19,20^. In addition, iPSC models have been used to study aberrant development in several neurodevelopmental disorders^21,22^. The ubiquity of stem cell differentiation protocols provides a unique opportunity to study the self-assembly of neural networks in a dish.

In this report, we introduce a method to study network features of developing human neural networks at the global and single-cell levels. We use long-term imaging coupled with automated image analysis to develop network representations of cell spatial topology and assign spatial coordinates to individual cells (**Figure 1**). We use our method to demonstrate that two independent human NPC cell lines exhibit a similar rise and fall in spatial network efficiency that characterizes the maturation of *in vitro* neural networks. We demonstrate that high spatial network efficiencies at intermediate stages of neural differentiation are linked with high levels of spontaneous electrical activity. We also use the graph method to uncover the spatial coordinates of specific cell types in developing cultures and aberrant spatial phenotypes in a neurodevelopmental disease model. The paradigm presented here can be used to uncover fundamental features of neural network formation from progenitor cells and link cellular spatial organization to function.

**Figure 1.**
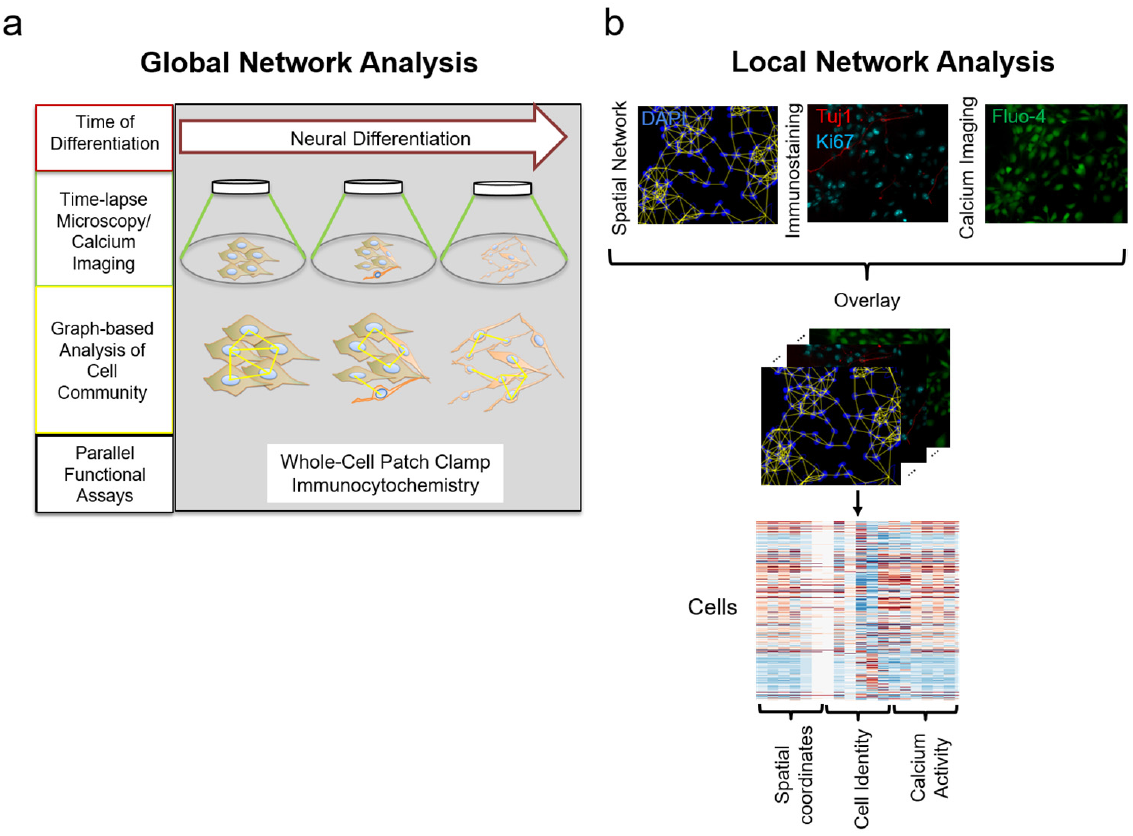
Schematic of the Living Neural Networks paradigm. **(a)** Global network analysis to uncover broad trends in NPC spatial arrangement, with parallel functional assays. **(b)** Local network analysis to reveal spatial coordinates of individual cells in culture.

## RESULTS

### Functional characterization and spatial network representation of differentiating NPC cultures

In the first part of this study, we used primary hNP1 neural progenitor cells derived from H9 human embryonic stem cells. These cells were maintained as undifferentiated, mitotic progenitor cells in the presence of basic fibroblast growth factor (bFGF). Withdrawal of growth factors from culture medium was used to induce spontaneous differentiation of hNPCs^23^.

First, we performed immunocytochemistry and whole-cell patch clamp electrophysiology experiments to uncover the time course of functional development in differentiating hNP1 cells. Multipotent NPCs prior to beginning neural induction were uniformly positive for Nestin, a Type VI intermediate filament protein expressed by dividing neural progenitor cells (**Figure 2a**). Cells at day 14 of neural induction were positive for microtubule-associated protein-2 (MAP2), a protein associated with dendrite formation in maturing neurons (**Figure 2b**). Analysis of peak inward and outward currents from voltage-clamp experiments showed that cells at all developmental stages exhibited equivalent levels of outward currents, but showed increasing magnitudes of inward currents (**Figure 2c, d**). Inward currents are typically driven by voltage-gated sodium channels and their presence indicates a more mature neuronal phenotype. Furthermore, weak action potentials could be elicited from cells showing inward currents at later time points (3/11 cells at day 14) through current injection (**Figure 2e**). These experiments demonstrated that Nestin-positive hNP1 cells matured over 14 days to MAP2-positive neurons, with neuronal fate commitment likely occurring between days 4-8, as indicated by the appearance of functional neuronal phenotypes in that time period.

**Figure 2.**
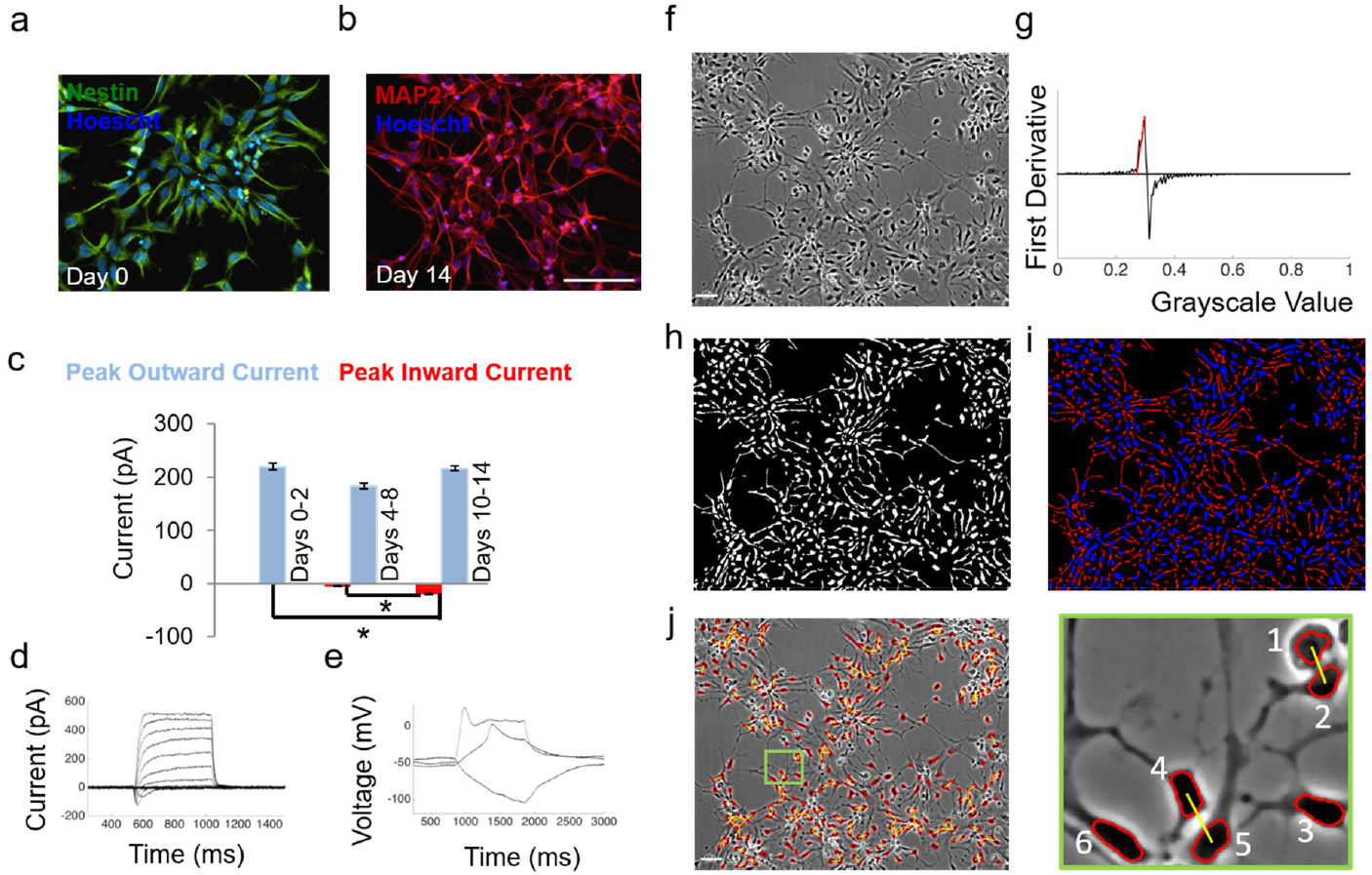
Functional characterization and spatial network representation of differentiating NPCs. **(a)** hNP1 cells at day 0 stain positively for Nestin. **(b)** Cells at day 14 stain positive for MAP2. In (a-b) nuclei are labeled by Hoescht; scale bar = 100μm. **(c)** Peak inward and outward currents determined through whole-cell patch clamp electrophysiology. Sample sizes: n=17, n=25, n=33 cells recorded for day 0-2, day 4-8 and day 10-14 respectively. Error bars represent SEM; *p < 0.05 from two-sample t-test. **(d)** Voltage-gated inward and outward currents seen in a cell at day 14. Voltage steps applied were from −60mV to +90mV in 10mV increments. **(e)** Weak action potentials evoked from the same cell through current injection. Magnitudes of current injected are −30pA, +20pA and +120pA from holding. **(f)** Representative phase contrast image of hNP1 cells, shown at day 3; scale bar = 50 μm. **(g)** First derivative of the pixel intensity histogram, with a linear fit to the ascending portion shown as a red line. The point where this line met the x-axis was used as a threshold for segmentation. **(h)** Binary image obtained upon thresholding the grayscale image. **(i)** Separation of linear features through morphological opening of the binary image yields cell bodies (blue) and neurites (red). **(j)** Phase contrast image from (f) with soma boundaries overlaid in red, and proximity edges shown in yellow. Inset shows six soma, of which two pairs (1, 2) and (4, 5) are connected by proximity edges; the intercellular distance for these two pairs are smaller than their average diameter multiplied by a scaling factor S = 2; Soma 3 and 6 are isolated nodes since they are not sufficiently close to any other soma. All microscope images are displayed with enhanced contrast for easy visualization.

In order to uncover topological changes in differentiating hNP1 cells, we combined long-term imaging of differentiating cultures with a graph-based approach for quantifying cell community structure. Differentiating cultures were imaged at days 0, 3, 6, 9, 12 and 14 after withdrawal of bFGF. An additional dataset was obtained by imaging differentiating cultures at 1-hour intervals for a total duration of 8 days (**Supplementary Video 1**). Selected image sequences were analyzed using custom image-processing algorithms, resulting in the extraction of soma and neurites for each phase-contrast image (**Figure 2f-i**) (see Methods for details).

We built network representations of spatial topology by denoting cell soma as nodes and using spatial proximity between soma to assign edges (**Figure 2j, k**). The resulting adjacency matrix, *A*, represented the spatial topology of cells, where *A_i,j_* = 1 if an edge existed between cells *i* and *j*, and 0 otherwise. In this manner, we constructed non-weighted, undirected graphs representing hNP1 communities from microscope images (**Supplementary Video 2**).

### Structure and information flow in NPC spatial graphs

In order to describe the structure and topology of hNP1 spatial graphs, we evaluated 17 metrics derived from graph theory that were computed and normalized appropriately to account for network size (**Table 1**)^24^. The network metrics provide information on various aspects of the graph structure such as information flow, connectivity and abundance of motifs (repeating patterns of cell arrangements). Through analyses of the covariance matrix of 17 metrics via hierarchical clustering, we were able to identify several strong positive correlations among degree-related metrics including average degree, average neighbor degree and degree variances (**Figure 3a**). We also identified negative metric correlations including those between network efficiency and number of connected components, as well as between clustering coefficient and all degree-related metrics. In the following section, we focus on metrics that have intuitive biological interpretations, their trends across time of differentiation, and their observed relationships with other metrics. Additional trends in metrics are presented in **Supplementary Figure 4**.

**Figure 3.**
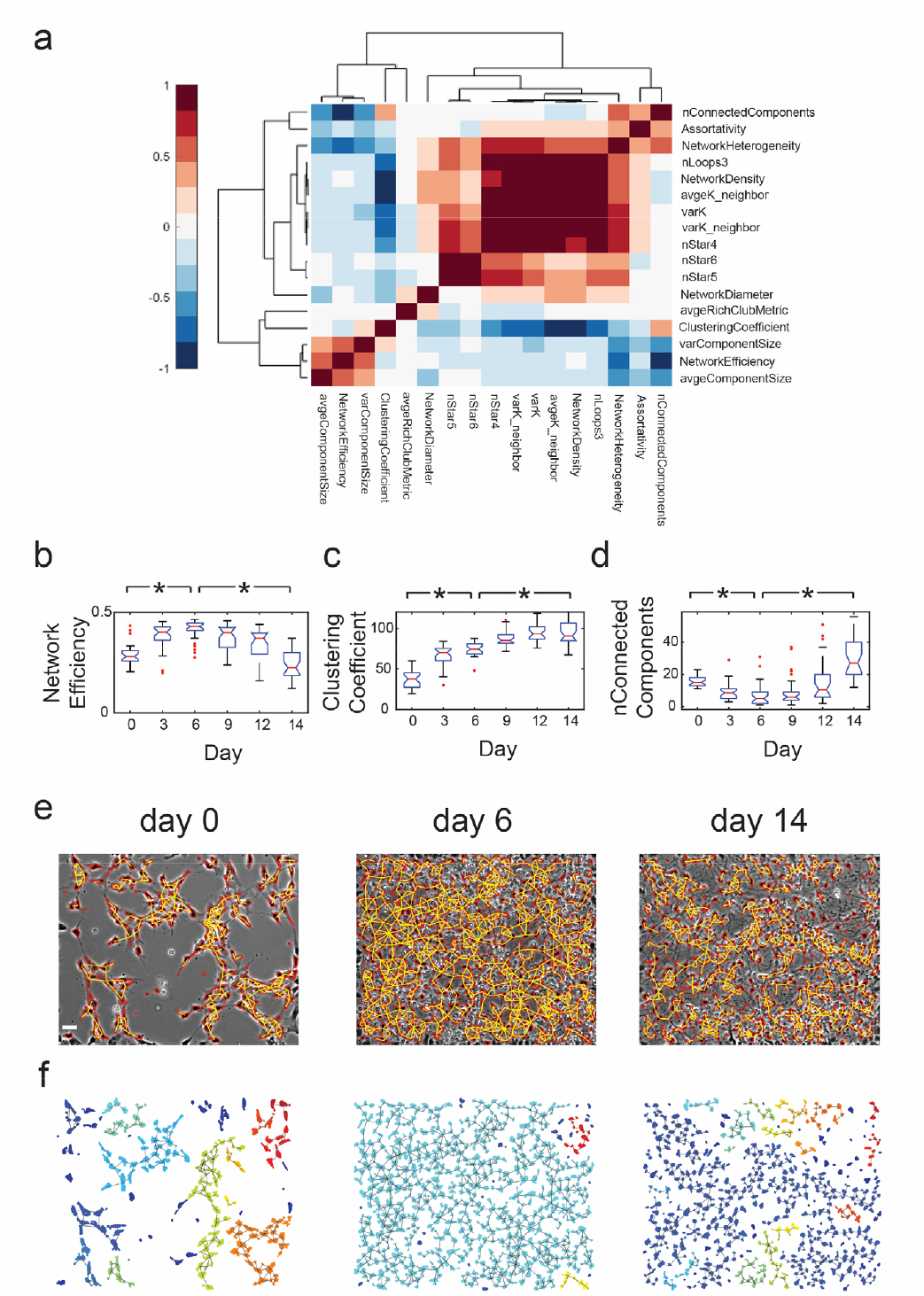
Spatial network efficiency is highest at intermediate stages of differentiation. **(a)** Correlation heatmap of all metrics obtained by hierarchical clustering of the covariance matrix. Clustering was performed using average linkage and Pearson correlation as the distance metric (shown in legend). Rows and columns are labeled with shorthand for metrics (Table 1). **(b)** Box plot of network efficiency across time. **(c)** Box plot of clustering coefficient across time. **(d)** Box plot of number of connected components across time. **(e)** Spatial graph representations of images taken at day 0, day 6 and day 14. Cell soma are outlined in red and edges are shown in yellow; scale bar = 50μm. **(f)** Cell soma from the images in (e), with each connected component labeled with distinct colors. For box plots in (b-d), red notches represent median (*Q*2), length of boxes represent interquartile range (*IQR*), length of notches represent 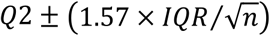, whiskers represent *Q*1 − (1.5 × *IQR*) and *Q*3 + (1.5 × *IQR*) and red circles represent outliers. *Q*1 = 25^th^ percentile, *Q*2 = median, *Q*3 = 75^th^ percentile, *n* = 30 data points for each box plot. * p < 0.00029 from Wilcoxon signed rank test (significance threshold adjusted using Bonferroni correction for 17 statistical tests to 0.005/17 = 0.00029)

**Table 1.** Global metrics computed, their descriptions, and mode of normalization to account for the network size. n = number of nodes, m = number of edges.

| Graph Metrics | Shorthand | Definition | Normalization |
| --- | --- | --- | --- |
| Network Density | NetworkDensity | Average degree of graph, normalized by total maximum possible degree<br>$\text{NetworkDensity} = \frac{2m}{n(n-1)}$ | Maximum possible degree, $(n-1)$ |
| Variance in Degree | varK | Variance of normalized node degree sequence | Node degree sequence normalized by maximum possible degree, $(n-1)$ |
| Network Heterogeneity | NetworkHeterogeneity | Standard deviation of normalized node degree sequence divided by mean of degree sequence | - |
| Average Neighbor Degree | avgeK_neighbor | Average degree of node neighborhood, across all nodes | Maximum possible degree, $(n-1)$ |
| Variance in Neighbor Degree | varK_neighbor | Variance of the normalized average neighbor degree sequence | - |
| Network Efficiency | NetworkEfficiency | The average reciprocal of shortest path length across all pairs of nodes, $E$ | Average network efficiency of 100 random graphs generated through degree-preserving rewiring, $E_{rand}$. Random graph generation is illustrated in **Supplementary Figure 3.** |
| Average Clustering Coefficient | ClusteringCoefficient | Fraction of total possible links among the neighbors of a node that are actually present, averaged across all nodes, $C$ | Average clustering coefficient of 100 random graphs generated through degree-preserving rewiring, $C_{rand}$ |
| Number of connected components | nConnectedComponents | Number of disconnected sub-graphs in main graph | - |
| Average Size of Connected Components | AvgeComponentSize | Average number of nodes in each connected component | Total nodes, $n$ |
| **Variance in size of connected components** | VarComponentSize | - | - |
| **Network Diameter** | NetworkDiameter | Longest shortest path length of network | Longest possible path, (n-1) |
| **Triangular loop count** | nLoops3 | Number of loops of 3 nodes | Total possible number of triplets, ${}^nC_3$ |
| **4-star motif Count** | nStar4 | Number of star motifs with one hub and three spokes | Total possible number of 4-tuples, ${}^nC_4$ |
| **5-star motif count** | nStar5 | Number of star motifs with one hub and four spokes | Total possible number of 5-tuples, ${}^nC_5$ |
| **6-star motif count** | nStar6 | Number of star motifs with one hub and five spokes | Total possible number of 6-tuples, ${}^nC_6$ |
| **Rich-Club Metric Average** | AvgeRichClubMetric | Measure of the tendency of nodes with high number of links to be well connected among each other<sup>26</sup>; Computed for threshold degrees between 1 and (n-1) | Average Rich-Club Metric of 100 random graphs generated through degree-preserving rewiring, $RCM_{rand}$ |
| **Assortativity** | Assortativity | Pearson correlation coefficient of degrees between pairs of linked nodes<sup>27</sup>. | - |

Network efficiency and clustering coefficient are commonly used measures of efficiency in global and local information flow^25^ (**Table 1**). When applied to hNP1 networks, these metrics describe the efficiency of information exchange at the network-wide and local neighborhood levels through cell soma proximity (compared to random graphs obtained through degree-preserving rewiring) (**Supplementary Figure 3**). In this context, information exchange could include the flow of ions through gap junctions or the diffusion of chemical signals from cell to cell. Evaluation of these metrics in hNP1 networks sampled across 30 different spatial locations from two biologically independent experiments showed that network efficiency increased from day 0 to day 6 and then decreased from day 6 to 14, while clustering coefficient rose constantly from day 0 to 14 (**Figure 3b,c**). Thus, there appears to be a transition from topologies favoring global information flow to those favoring a hierarchical form of communication, occurring between day 6 and 14 of differentiation.

The metric correlation heatmap showed a strong negative correlation between network efficiency and number of connected components in the graph (**Figure 3a**). The number of connected components is a count of the number of disconnected sub-graphs in the main network and is a measure of the connectivity of the graph – a graph with a high number of connected components has a low connectivity (**Figure 3d**). NPC networks at day 0, 6, and 14 are shown in **Figure 3e** and the corresponding connected components are shown in **Figure 3f**. Analysis of NPC networks at day 0, 6, and 14 identified the formation of a giant connected component likely due to continued cell proliferation through day 6 of differentiation (**Figure 3e,f**). This increase in the connectivity of the network resulted in an increase in network efficiency (**Figure 3b**). The subsequent disaggregation of the large component into smaller modules between days 6 to 14 contributes to the decrease in network efficiency seen at later developmental stages (**Figure 3b**).

### Development of functional and spatial networks

We next probed the relationship between functional and spatial networks in developing NPCs using ReNcell VM immortalized human neural progenitor cells. Differentiation induced by growth factor withdrawal led to the formation of dense networks within 5 days, rapid exit from the cell cycle (as seen by reduced expression of Ki67) and formation of β(III)-tubulin-positive neurons. (**Supplementary Figure 6a**).

We performed calcium imaging using the fluorescent calcium indicator Fluo-4 to record spontaneous activity in differentiating ReNcell VM cultures at days 1, 3 and 5, and employed cross-correlation analysis to infer functional connectivity in the networks (**Figure 4a, Supplementary Figure 7**). Analysis of functional networks revealed that cultures at day 3 had significantly more activity than those at days 1 and 5, as measured by the fraction of active cells (**Figure 4b, Supplementary Videos 3-6**). Interestingly, the functional network was not restricted to cells with short intercellular distances, with cells in the whole field of view (832μm x 702μm) having highly correlated calcium activity (**Figure 4c**). Further, cultures at day 3 displayed waves of calcium activity propagating through many neighboring cells, that were not seen at later time points (**Supplementary Video 5**). When maintained in proliferation medium, ReNcell VM cultures continued to divide and did not differentiate into neurons (**Supplementary Figure 6b**). Further, the level of spontaneous activity in proliferation medium remained constant through day 5 of measurement, indicating that the trends in functional network activity seen in differentiating cultures was unique to the formation of neural networks (**Supplementary Figure 6c**).

**Figure 4.**
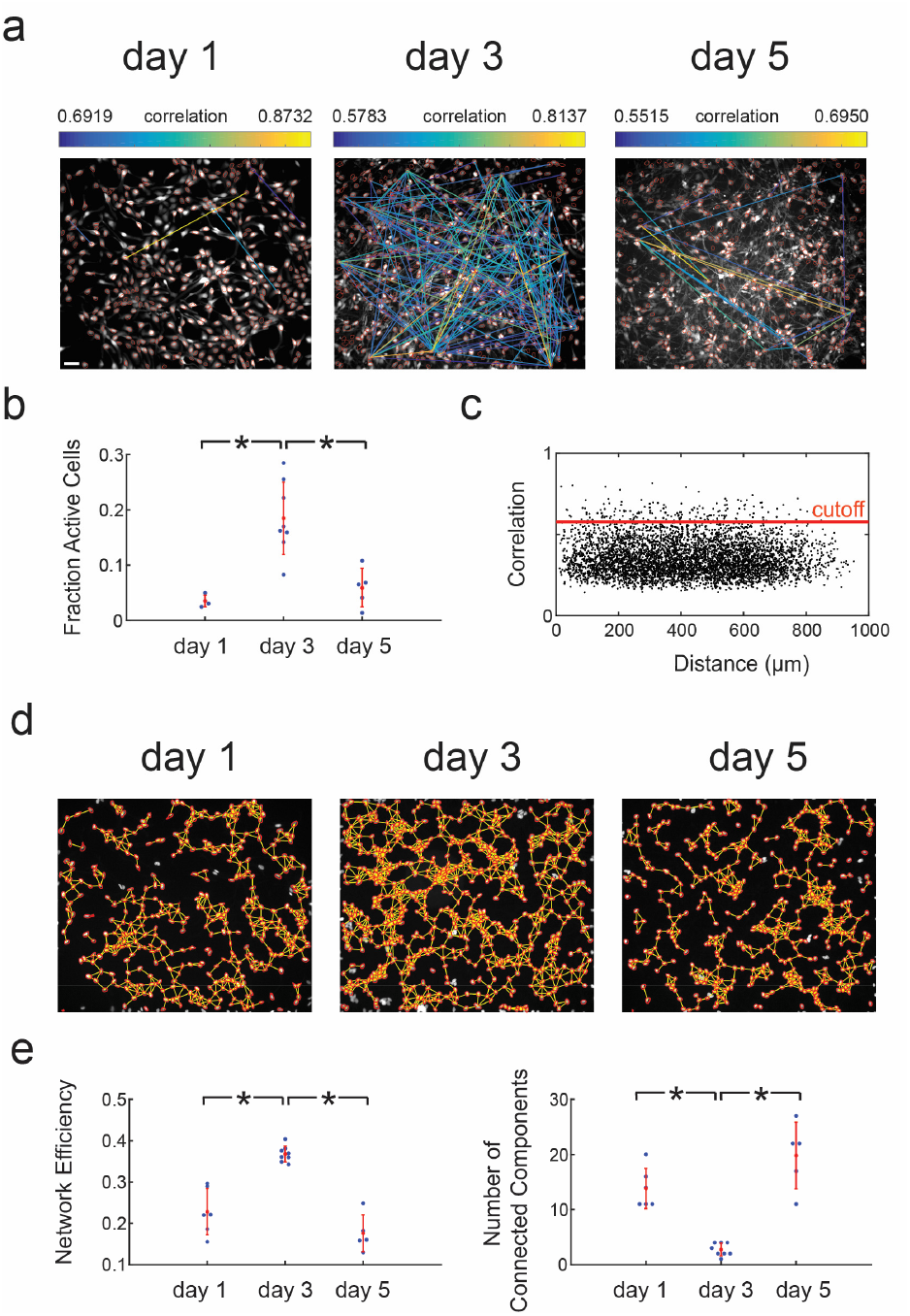
Functional and spatial networks in ReNcell VM NPCs. **(a)** Functional networks obtained through calcium imaging with Fluo-4 in developing NPC networks at days 1, 3 and 5. Correlations between cells are shown as a network plot overlaid on the maximum intensity image from calcium image sequences; scale bar = 50μm. **(b)** Fraction of active cells in the network. Active cells are defined as cells whose normalized fluorescence traces have three or more calcium transients; *p < 0.005 from two-sample t-test. **(c)** Plot of correlation versus intercellular distance for day 3 network shown in (a). Correlation threshold generated from shuffled dataset is shown as a red line. **(d)** Spatial networks overlaid on immunofluorescence images of nuclei stained with Hoescht dye; scaling factor = 3. The nucleus images correspond to the images shown in (a). **(e)** Network efficiency of spatial networks peaks at day 3. Number of connected components shows the inverse trend. Sample sizes: Day 1 (n=5); Day 3 (n=8); Day 5 (n=5) for all plots. Red notches show mean and standard deviation; *p < 0.00029 from two-sample t-test (significance threshold adjusted using Bonferroni correction for 17 statistical tests to 0.005/17 = 0.00029). All microscope images are displayed with enhanced contrast for easy visualization.

We next built spatial graphs using nucleus images from the same cultures in which calcium imaging was performed (**Figure 4d**). Spatial networks were most efficient at day 3 with the fewest number of connected components (**Figure 4e**). The rise and fall of network efficiency in ReNcell VM networks mirrored the trends seen in hNP1 networks, with the time course of the trends indicative of network maturation (**Supplementary Video 7**). Further, the peak in spatial network efficiency coincided with the most active functional networks. This leads us to conclude that the high spatial efficiency of NPC networks at intermediate time points of differentiation facilitates high levels of network-wide spontaneous activity.

### Single-cell analysis of developing neural networks

To identify the spatial and functional roles of different cell types in developing neural networks, we used neural stem cells derived from the NCRM-5 human iPSC line^28^. NCRM-5 NSC cultures differentiated into dense networks of neurons over a period of 28 days (**Supplementary Figure 8**).

We next performed immunostaining for proliferating cells (Ki67) and new neurons (β(III)-tubulin/Tuj1) at day 3 of differentiation and leveraged the graph theoretic approach to evaluate spatial features of individual cells (**Table 2**). This analysis revealed that Ki67+ proliferating cells had a high degree or number of neighbors compared to Ki67- non-proliferating cells in NCRM-5 cultures (**Figure 5a**). Further, Tuj1+ neurons had a high clustering coefficient compared to Tuj1- cells in differentiating NCRM-5 NSCs (**Figure 5b-d**). Cells with high clustering coefficient are likely to be part of cliques, a common feature of cells at the edge of clusters due to geometric constraints (**Figure 5d**). Thus, our results suggest that proliferating cells tend to be close to the center of clusters where they are surrounded by many neighbors, while newly born neurons are found mostly at the edge of clusters.

**Figure 5.**
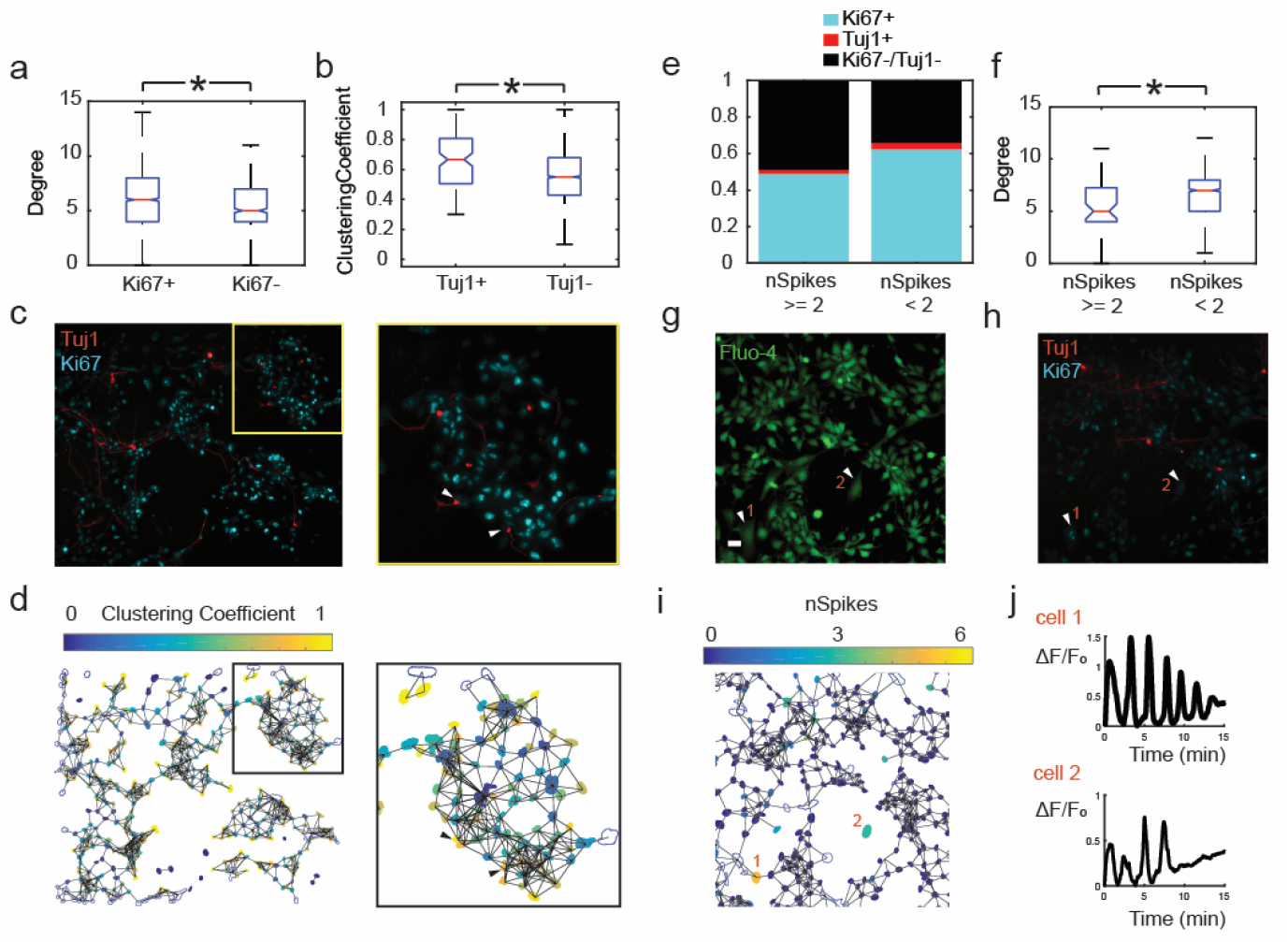
Multiparametric single-cell analysis of day 3 NCRM-5 NSC cultures. **(a)** Boxplot of degree versus Ki67 status. Ki67+ cells have higher degree than Ki67- cells. **(b)** Boxplot of clustering coefficient versus Tuj1 status. Tuj1+ cells have higher clustering coefficient than Tuj1- cells. **(c)** Immunofluorescence image of day 3 NCRM-5 NSC culture with Tuj1 and Ki67 stains; scale bar - 50μm. Inset is shown in yellow box with arrows pointing to Tuj1+ neurons at the network periphery. **(d)** Clustering coefficient of individual cells shown as a heatmap with spatial network overlaid; scaling factor = 3. Nucleus (DAPI) images corresponding to (c) were used to create the spatial graph. Inset shows the same cells as inset in (c). Arrows point to Tuj1+ neurons at the network periphery with high clustering coefficient. **(e)** Proportions of cell types comprising high-spiking versus low-spiking calcium imaging. **(f)** Boxplot of degree versus spiking characteristics. High-spiking cells have lower degree than low-spiking cells. **(g)** Frame from calcium imaging sequence for day 3 NCRM-5 NSC culture; scale bar - 25μm. Arrows point to high-spiking, morphologically distinct cells with few neighbors. **(h)** Immunofluorescence image corresponding to (g) identifying Tuj1+ and Ki67+ cell types. **(i)** Heatmap of number of spikes of spontaneous calcium activity within a representative 15 min imaging window with spatial network overlaid. **(j)** Calcium traces from two high-spiking cells. *p < 0.0071 from two-sample t-test (significance threshold adjusted using Bonferroni correction for 7 statistical tests to 0.05/7 = 0. 0071).

**Table 2.** Metrics describing local network features, calculated at the level of individual cells.

| Graph Metrics | Symbol | Definition |
| --- | --- | --- |
| Degree | $k$ | Number of neighbors (one link away) |
| Average Neighbor Degree | $k_n$ | Average degree of all neighboring cells |
| Clustering Coefficient | $C$ | Fraction of total possible links among the neighbors of a node that are actually present, averaged across all neighbors |
| Local Efficiency | $E_l$ | Average shortest path length in local neighborhood |
| Node Closeness Centrality | $c_n$ | Sum of reciprocal distances to all other nodes |
| Node Betweenness Centrality | $w_n$ | Number of shortest paths that pass through the node |

In order to investigate the functional role of individual cells, we performed calcium imaging using Fluo-4 on day 3 cultures, followed by immunostaining and co-registration of the immunostain image with the calcium video (**Figure 1b, Supplementary Video 8-10**). This analysis revealed that high-spiking cells had a greater proportion of Tuj1−/Ki67− cells and a lower degree than low-spiking cells (**Figure 5e,f**). Through visual inspection, this high-spiking population of cells exhibited qualitatively larger morphologies (**Figure 5g-j**).

### Analysis of a neurodevelopmental disease model reveals aberrant network features

To validate our network model for applications to study diseases, we performed spatial network analyses using an iPSC disease model of Smith-Lemli-Opitz syndrome (SLOS), an autosomal recessive developmental disorder resulting from mutations in *DHCR7* which produces pronounced neurological deficits^29^. Previous studies have shown accelerated differentiation of neural progenitors derived from patients with SLOS^22^, likely caused by decreased activity of the canonical Wnt/β-catenin signaling pathway in these cells.

Here, we compared the global and individual cell spatial features of day 3 differentiating cultures of control (NCRM-5) and SLOS iPSC-derived NSCs (CWI 4F2). At the global level, CWI 4F2 cultures were more homogeneous than NCRM-5 cultures, as indicated by the presence of fewer connected components and lower network heterogeneity (**Figure 6a-d, Table 1**).

**Figure 6.**
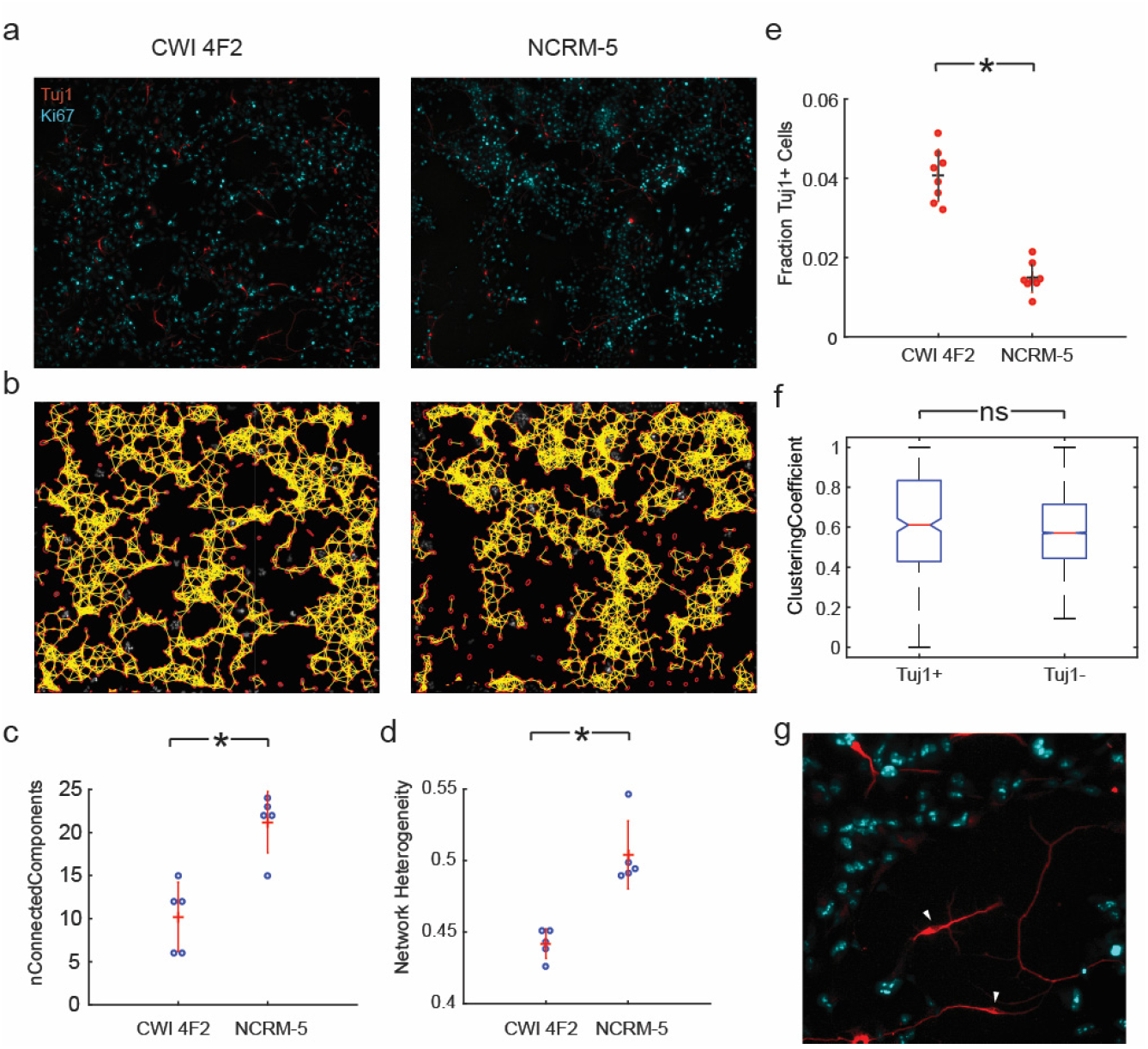
Spatial network analysis of iPSC-derived NSCs reveals deficits in network connectivity in a neurodevelopmental disorder model. **(a)** Immunostained images from day 3 cultures of CWI 4F2 SLOS NSCs and control NCRM-5 NSCs; scale bar = 50 μm. **(b)** Nuclei stained by DAPI corresponding to images in (a), with spatial graph overlay; scaling factor = 3. **(c)** Number of connected components in NCRM-5 cultures is higher than CWI 4F2. **(d)** Network heterogeneity of NCRM −5 cultures is higher than CWI 4F2; *p < 0.0029 from two-sample t-test (significance threshold adjusted using Bonferroni correction for 17 statistical tests to 0.05/17 = 0. 0029). **(e)** CWI 4F2 NSCs exhibit accelerated differentiation into Tuj1+ neurons compared to NCRM-5 NSCs. **(f)** Clustering coefficient of Tuj1+ cells shows no difference to that of Tuj1− cells in day 3 CWI 4F2 cultures. **(g)** Portion of differentiating CWI 4F2 NSCs immunostained image in (a), with arrows pointing to disconnected Tuj1+ neurons contributing to low clustering coefficient in CWI 4F2 cultures.

Our analyses both confirmed accelerated neural specification in CWI 4F2 cultures through day 3 and revealed that Tuj1+ neurons in day 3 CWI 4F2 cultures did not have a high clustering coefficient compared to Tuj1-cells, as was the case in NCRM-5 cultures (**Figure 6e, f**). This was due to the presence of many more ‘lone’ neurons with higher neurite extensions in the CWI 4F2 cultures (**Figure 6g**). Thus, network analysis revealed the presence of global and local features of spatial organization in neural cell types in a neurodevelopmental disorder, validating the utility of the Living Neural Network model for the study of neurological diseases.

## DISCUSSION

Topological and functional analysis of *in vitro* neural networks has the potential to uncover basic organizational principles of their *in vivo* counterparts. Our study provides a new approach which leverages the directed differentiation of human stem cells to study the self-assembly of *in vitro* neural networks from neural progenitor cells. We quantified the spatial organization of immature neural cells during differentiation, using a unique application of graph theory. The experimental paradigm presented here enabled us to uncover relationships between spatial topology of NPC communities and functional maturation of developing neural circuits, and allowed us to develop hypotheses about the role of certain topologies on NPC function.

Because chemical and electrical signaling between neural progenitor cells often involves direct contact between adjacent cells, we expect spatial cell organization to be an important aspect of neural development. Two examples where spatial organization would play a role include Notch/Delta signaling and gap junctional communication between cells. The Notch/Delta signaling pathway, which influences the proliferation and neuronal fate commitment of progenitor cells^30,31^, is an example of juxtacrine chemical signaling. The canonical Notch signaling pathway functions through the binding of a transmembrane ligand on one cell with the transmembrane receptor on a contacting cell, resulting in the release of the notch intracellular domain (NICD) to initiate downstream signaling cascades in the contacting cell^32^. In addition, immature neural circuits are known to display spontaneous electrical activity, which is an important aspect of their proper development^13,14^. Gap junctions or electrical synapses allow direct access between cells and result in exchange of ions and growth factors, and these are known to be important in the propagation of spontaneous electrical activity. Indeed, neural progenitor cells have been shown to display structured and synchronous calcium activity, which depends on gap junctions and which promotes cell proliferation^12^. More broadly, structured cell-cell communication has been implicated in coordinated chemosensing^33^ and migration during development^34^. Thus, the methods of cell-cell communication employed by immature neural cells indicate the significance of spatially organized electrical and chemical signaling.

We found that the rise and fall in spatial network efficiency was a reproducible feature in NPC networks, and believe this is a characteristic feature of the transition from immature NPC networks to mature neuronal networks. Global network efficiency is low in proliferating NPC topologies, rises to a peak in intermediate cultures, and then drops off as cells mature into neuronal networks. The trend in network efficiency is intuitively explained by its negative correlation with the number of connected components, a measure of cell connectivity. Cell proliferation in the early stages of differentiation causes the merging of many disconnected clusters of cells into a giant connected component, which leads to a rise in the overall spatial connectivity and reduction in the average path length. The reorganization of the giant component into smaller modules at later stages leads to a reduction in network efficiency.

Of particular developmental importance, we determined the peak of spatial network efficiency coincides with the appearance of electrophysiologically mature neurons in culture and high levels of spontaneous network-wide calcium activity, respectively, in the cell lines we studied (hNP1 and ReNcell VM cells). Additionally, levels of spontaneous activity drop off in more mature cultures which have significant neurite outgrowth and more clustered cell bodies. The spatial and functional architecture of mature cultures is consistent with previous evidence of highly clustered units developing in neuronal cultures^5–7^. The reduction in network-wide spontaneous activity in more mature cultures is consistent with a transition from a global to a hierarchical structure of communication.

By adapting the graph-based approach at the single-cell level, we also found that specific cell types have unique spatial coordinates in developing NSC cultures derived from human iPSCs. We found that Ki67+ proliferating cells had a higher average number of neighbors than non-proliferating cells and newly born Tuj1+ neurons had high clustering coefficients indicative of their locations at the culture periphery. These results correspond to previously observed features of polarized neuroepithelium – prevalence of mitosis near the lumen and migration of differentiated cells to the periphery^19^. Further, by coupling calcium imaging experiments with spatial analysis of specific cell types, we found that high-spiking cells tend to have reduced numbers of neighbors and have a higher proportion of Tuj1-/Ki67- cells. Through visual inspection, we observed these cells had large morphologies. Based on their morphological and spatial properties, we hypothesize this cell population may represent a basal progenitor cell population previously observed in differentiating NSC cultures^19,35^.

We then leveraged spatial network analysis to uncover features of aberrant neural networks in an iPSC model of a neurodevelopmental disorder, Smith-Lemli-Opitz syndrome. Global network analysis demonstrated that developing SLOS cultures were more homogeneous than control NCRM-5 cultures at the same stage of differentiation. We believe this indicates that normal differentiation is characterized by the presence of NSC clusters of a wide size distribution, a feature that is disrupted in SLOS. Further, the higher number of neurons in SLOS cultures comprised a large number of ‘lone’ cells with high neurite extension. While the mechanisms underlying altered neural networks in SLOS are unclear, published findings related to cytoskeletal remodeling or diminished β-catenin signaling affecting cadherin function are possibilities^22,36^.

In this study, we show that network analysis provides unique information about the structure of neural progenitor cell communities at the local and global levels. It remains to be seen whether spatial topology of developing cultures is predictive of synaptic connectivity in mature neuronal networks. Several *in vivo* studies have provided evidence for a structure-functional relationship between adult neuronal wiring and the spatiotemporal origin of the constituent neurons. For example, sister excitatory neurons in the neocortex are more likely to develop synapses with each other rather than with other cells^37^, and the electrophysiological phenotypes of GABAergic interneurons have been shown to depend on the time and place of their birth^38^. Thus, the analysis of spatial topology in developing neuronal circuits in a controlled setting has the potential to uncover structure-function relationships in the resulting mature neural circuits.

The present study also lays the foundation for analysis of the role of cellular neighborhood on cell fate determination of individual progenitor cells. The expression of cell-fate determination factors such as bHLH transcription factors like Hes1 and Ngn2, and proteins involved in cell-cell communication pathways such as Notch/Delta proteins, have been shown to be tightly coupled with each other^17,39^. Computational modeling studies have predicted that Notch-Hes1 intercellular signaling affects differentiation and cell cycle progression of individual cells and this signaling is important for the maintenance of an optimal balance between differentiating cells and self-renewing progenitor cells^40^. The spatial dynamics of cell-cell signaling and its impact on single-cell differentiation status is an intriguing subject for future study.

In conclusion, we present a multiplexed approach integrating long-term imaging, automated image analysis, and graph theory to quantify the spatial and functional networks of neural progenitors during neural differentiation. The Living Neural Networks method introduces a tangible means to test theories about different forms of neural cell communication and their role in shaping functional neural networks. Insights from this study help further our understanding of the fundamental design features of the brain.

## METHODS

### Cell culture

Human neural progenitor cells (hNP1) derived from H9 human embryonic stem cells were obtained from ArunA Biomedicals (Athens, GA). Cells were expanded on tissue culture flasks pre-coated with either fibronectin (Sigma-Aldrich) or Matrigel (BD Biosciences), in proliferation medium consisting of AB2 basal neural medium, ANS neural supplement (both supplied by manufacturer), 10 ng/ml leukemia inhibitory factor (LIF; EMD Millipore), 20 ng/ml basic fibroblast growth factor (bFGF; R&D Systems), 2 mM GlutaMAX supplement (Life Technologies) and penicillin/streptomycin (Life Technologies). For neural differentiation experiments, cells were cultured in medium lacking bFGF.

ReNcell VM immortalized human neural progenitor cells derived from the ventral mesencephalon of human fetal brain were purchased from EMD Millipore. Cells were expanded on tissue culture flasks coated with laminin (Life Technologies), in media containing DMEM/F12 (Life Technologies), supplemented with B27 (Life Technologies), 2 μg/ml heparin (StemCell Technologies), 20 ng/ml bFGF (EMD Millipore), 20 ng/ml EGF (Sigma) and penicillin/streptomycin (Life Technologies). For differentiation experiments, cells were cultured in medium lacking bFGF and EGF.

Neural stem cells were derived from two human iPSC lines, NCRM-5 and CWI 4F2, following published protocols^22^ (**Supplementary Table 3**). NSCs were cultured on dishes coated with Poly-L-Ornithine and Laminin, in media containing DMEM/F12 (Life Technologies), B27 without vitamin A (Life Technologies), 20 ng/mL EGF (Sigma), 20 ng/mL bFGF (Stemgent) and penicillin/streptomycin (Life Technologies). NSCs were passaged using Accutase (Life Technologies) in medium containing 10 μm ROCK inhibitor (Y276332; Reagents Direct). Differentiation experiments were carried out in differentiation medium containing Neurobasal media (Life Technologies), B27 with Vitamin A (Life Technologies), 10 ng/mL BDNF (Peprotech), 10 ng/mL GDNF (Peprotech), GlutaMAX supplement (Life Technologies) and penicillin/streptomycin (Life Technologies). The NCRM-5 and CWI 4F2 iPSC lines were originally derived within the intramural program of the National Heart, Lung, and Blood Institute (NHLBI iPSC Core) and the Eunice Kennedy Shriver National Institute of Children’s Health and Human Development (laboratory of Forbes D. Porter).

### Electrophysiology

For whole-cell patch clamp experiments, cultures were maintained in extracellular recording solution containing 119 mM NaCl, 5 mM KCl, 10 mM HEPES, 2 mM CaCl_2_ and 1 mM MgCl_2_, titrated to a pH of 7.3. Pipettes (5-10 MΩ) were pulled from standard borosilicate glass capillaries and back filled with intracellular recording solution containing 8 mM NaCl, 10mM KCl, 5 mM HEPES, 0.06 mM CaCl_2_, 5 mM MgCl_2_, 130 mM potassium gluconate and 0.6mM EGTA, titrated to a pH of 7.4. Recordings were performed using a MultiClamp 700A amplifier and a Digidata 1550 Data Acquisition System coupled with Clampex 10.4 software (Molecular Devices). Traces were analyzed in MATLAB.

In voltage-clamp experiments, cells were held at a holding potential of −50 mV and given a series of voltage steps from −90 to +100 mV. In current-clamp experiments, cells were held at approximately −70 mV through minimal current injection before application of a series of current steps ranging from −40 to +120 pA. Magnitudes of the current steps were modified according to the input resistance. Peak outward current amplitude was measured 40 ms after the initiation of the voltage sweep. Peak inward current was defined as the maximum transient negative current at any command voltage.

### Immunocytochemistry

Cells plated on chambered cover glasses (Fisher Scientific) or glass coverslips were fixed in 4% PFA for 20 min, washed with PBS and incubated with blocking buffer containing 0.2% Triton-X (Sigma), 0.3M glycine and 10% goat serum (Jackson Immunoresearch Labs) for 1 hour. Cells were then incubated overnight in the following primary antibodies diluted in 10% goat serum: Mouse Nestin (1:200, Neuromics), Rabbit MAP2 (1:500, Millipore), Mouse Tuj1 (1:2000, Millipore), Rabbit Ki67 (1.43μL/mL, Abcam). Cells were rinsed and primary antibodies were detected using appropriate Alexa Fluor secondary antibodies. Nuclei were stained using either Hoescht or DAPI.

See **Supplementary Table 2** for a list of antibodies and dilutions.

### Image acquisition and segmentation

For live imaging experiments, hNP1 cells were plated at approximately 50% confluence on 12-well plates pre-coated with Matrigel and switched to differentiation medium 24 hours post-plating. Two datasets (biological replicates) were obtained by imaging the well plates at days 0, 3, 6, 9, 12 and 14 after withdrawal of bFGF from culture medium, using an automated stage Nikon Eclipse Ti-E Microscope. At the start of the experiment, five locations were chosen arbitrarily for each well, and the same locations were located and imaged at each time point. Imaging sessions lasted about 10 minutes and the plates were returned to the incubator after imaging. We also performed continuous imaging (**Supplementary Video 1**), for which the well plate was mounted on the stage of the microscope in a bold line cage incubator (Okolab) equipped with temperature control and gas flow rate control enabling a 37°C 5% CO_2_ environment. Images were acquired at 1-hour intervals for 8 days. In all time-lapse imaging experiments, 8-bit phase contrast images were acquired through a 10X objective (N.A. = 0.3) from a 1280 x 1080 pixel field of view using a Nikon DS-Qi1 camera. Physical pixel size was 0.64 μm.

Phase contrast image sequences were chosen for analysis based on the ability of a human observer to distinguish cellular features in the images. Images with large amounts of debris occluding cells were discarded manually. In this manner, a total of 16 and 14 image sequences (30 locations) for each of the 2 biologically independent datasets were chosen for analysis.

Selected grayscale images were pre-processed by applying a median filter with a neighborhood of 3x3 pixels to remove noise and segmented using an unbiased intensity-gradient thresholding approach^41^. Starting from the grayscale image, the first derivative of the pixel intensity histogram was calculated. Fitting a linear function to the ascending portion of the first derivative and extrapolating to the x-axis resulted in a grayscale threshold, which was used to generate a binary image distinguishing cellular features from the background. Morphological operations performed on the binary image were:

1. Small objects of size lesser than 50 pixels were removed to filter out noise and other imaging artifacts.
2. Morphological opening was performed using a disk structuring element of radius 4 pixels. This was done to separate linear features (neurites), and circular features (cell bodies).
3. Cell bodies were separated using connected component labeling using the default 8-connected neighborhood.
4. Cell body objects smaller than 150 pixels and those touching the image border were removed.

All parameters used in phase contrast image processing are listed in **Supplementary Table 1**.

In order to quantify the accuracy of our image processing algorithms, we compared the results with manual tracing of soma. These results showed a close agreement between the numbers of cells detected by our algorithm and by manual tracing at different time points (**Supplementary Figure 1**).

### Calcium imaging and analysis

Cells were plated on LabTek chambered cover glasses for calcium imaging experiments. Cells were loaded with culture medium containing 3 μM of the fluorescent calcium indicator Fluo-4/AM (Life Technologies) and Pluronic F-127 (0.2% w/v, Life Technologies) for 30 min at 37°C. Imaging of spontaneous calcium activity was performed at 37°C using a 20X objective lens (N.A. = 0.75), with 488 nm excitation provided through a SOLA SE Light Engine (Lumencor). 16-bit fluorescence images were acquired at a sampling frequency of 1 Hz for a total duration of 15 min, using a Zyla 5.5 sCMOS camera (Andor).

Following calcium imaging, samples were subjected to immunocytochemistry as described earlier. By navigating to the locations where calcium imaging was performed, manual co-registration was done to obtain immunofluorescence images for the same fields of view.

### Generation of functional networks from calcium imaging

Regions of interest (ROIs) were obtained by segmenting nucleus images using a local thresholding approach followed by the watershed algorithm. Undersegmented objects were algorithmically removed by discarding the top two percentile of object sizes obtained after segmentation.

Next, a time-varying fluorescence trace was calculated for each ROI. For each frame in the calcium fluorescence image stack, background (average pixel intensity of non-ROI regions in the image) was subtracted. Average fluorescence intensity for each ROI (*F*) was obtained by averaging pixel intensity values within the ROI for each time point. Baseline fluorescence (*F*_0_) for each ROI was calculated as the minimum intensity value in a window 90s before and after each time point. The normalized fluorescence trace for the ROI was then calculated as *F* − *F*_0_/*F*_0_. Cells with low activity were filtered out by discarding traces with less than three peaks and traces whose signal-to-noise ratio was lower than 1. Quality of the remaining traces was confirmed by manual inspection. This was done to avoid false positives in the cross-correlation analysis.

Functional networks were created following the method described by Smedler et al^42^, where cross-covariance between signals is used to assign functional connections between pairs of cells. A randomized dataset was generated by shuffling each signal in the original dataset at a random time point. The 99^th^ percentile of cross-covariance values for the randomized dataset was used as a threshold for determining significant correlations.

### Creation of spatial graphs

Spatial graphs were created from microscope images using cytoNet, software developed in-house^43^. For each pair of objects (soma/nuclei), a threshold distance for proximity was defined as the average of the two object diameters, multiplied by a scaling factor (S). If the Euclidean distance between the object centroids was lower than the threshold distance computed, then the pair of objects was connected with a “proximity edge” (**Figure 2j, k**). We chose a scaling factor of 2 for phase contrast images and 3 for nucleus immunofluorescence images based on similarity in network density for the resulting networks (**Supplementary Figure 2**).

Due to the high density of NCRM-5 cultures, quantification of global network metrics proved unfeasible (**Supplementary Figure 8**). However, qualitatively we observed the prevalence of highly clustered cell bodies at late stages of differentiation.

### Metric computation

All the network metrics described in Table 1 were computed using cytoNet. It is to be noted that not all metrics derived from graph theory have a ready biological interpretation, especially in the context of spatial graphs. For example, interpretation of metrics like degree-degree correlations and rich-club metric (**Supplementary Figure 4**) are limited, due to the implicit limit in the type of connections that are possible in spatial graphs. Keeping this in mind, we focused on analyzing metrics with an intuitive biological interpretation, i.e., information flow and connectivity.

Random graphs were constructed through degree-preserving rewiring, maintaining the degree distribution of the original graph (**Supplementary Figure 3**). Each link (edge) belonging to any given node in the original graph was randomly re-assigned to a node that was chosen from all possible nodes with uniform probability. Metrics computed for random graphs were averaged across 100 different realizations of the random graphs. This mode of random graph generation was chosen to eliminate finite-size effects inherent in other models of random graphs such as Erdõs-Rényi random graphs. To ensure robustness of the network metrics, we tested varying fields of view for the images, and confirmed the trends remained consistent (**Supplementary Figure 5**).

### Single-cell analysis

Consolidated multi-parametric datasets were obtained by performing calcium imaging followed by immunocytochemistry (**Supplementary Videos 8-10**). Functional data obtained through calcium imaging (e.g., number of spikes) was combined with cell identity information obtained through immunostaining (e.g., Ki67, Tuj1 status), and spatial features extracted using nuclei as described earlier. Cells within 100 pixels of the border of the field of view were excluded from analysis to eliminate border effects. Occasionally, colonies of cells were washed away during the immunostaining process. In these cases, the calcium imaging channel was used to obtain an approximate mask of such cells in order to obtain a complete image set for spatial analysis. Additionally, large masks likely representing undersegmented objects were excluded from analysis.

### Code and data availability

All MATLAB code and data that support the findings of this study are available from the corresponding author upon request. The cytoNet user interface can be found at http://qutublab.rice.edu/cytoNet/

## ACKNOWLEDGEMENTS

We thank Dr. Byron Long, Dr. David Noren, Dr. André Schultz, Chenyue Wendy Hu and Dr. Ka Wai Lin for helpful discussions and comments on the manuscript, Hanyang Li and Kylie Balotin for assistance with manual tracing of cells, Daniel Murphy and Dr. Guillaume Duret for technical assistance. This work was supported by NSF Career Grant 1150645 to A.A.Q., NSF Neural and Cognitive Systems grant 1533708 to A.A.Q. and J.T.R., and NIH grant 5P20GM103620 to K.R.F. A.S.M. was supported through NSF IGERT training grant 1250104.

## AUTHOR CONTRIBUTIONS

All authors designed the experiments. A.S.M. performed the experiments. A.A.Q., A.S.M. and N.E.G. analyzed the data. All authors contributed to writing the manuscript. A.A.Q. and J.T.R. supervised the work. K.R.F supplied iPSC-derived NSC lines and supervised neural differentiation experiments.

## COMPETING FINANCIAL INTERESTS

The authors declare no competing financial interests.

## Supplementary Information

**Supplementary Figure 1.**
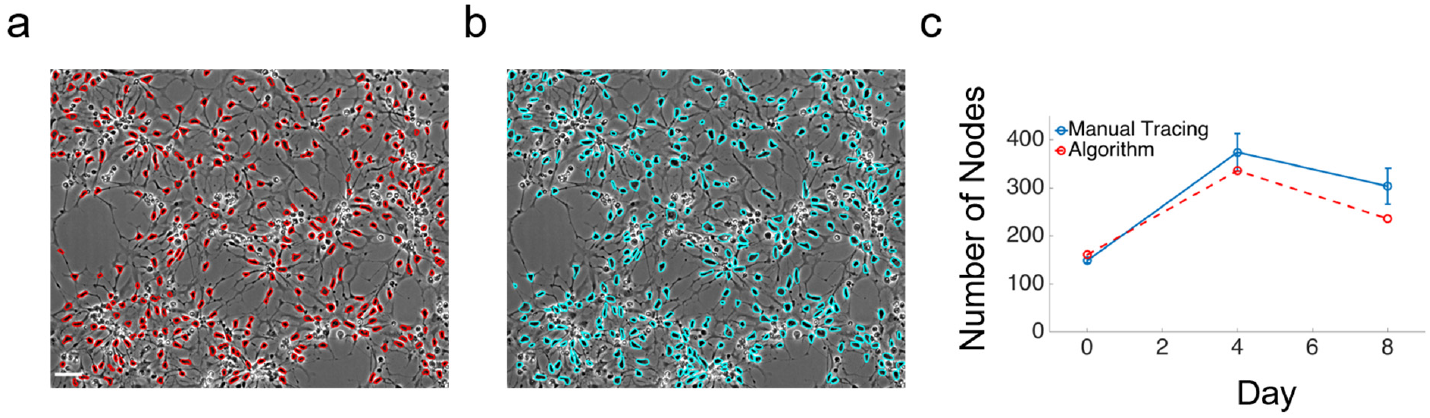
Comparison of automated image segmentation and manual tracing of cell soma. **(a)** hNP1 image at day 4 with cell soma outlines picked out by algorithm outlined in red; scale bar = 50 μm. **(b)** Same image in (a), with manual tracing of cell soma outlined in cyan. **(c)** Comparison of number of cell soma picked out by algorithm and manual tracing by 3 independent observers. Error bars indicate SEM.

**Supplementary Figure 2.**
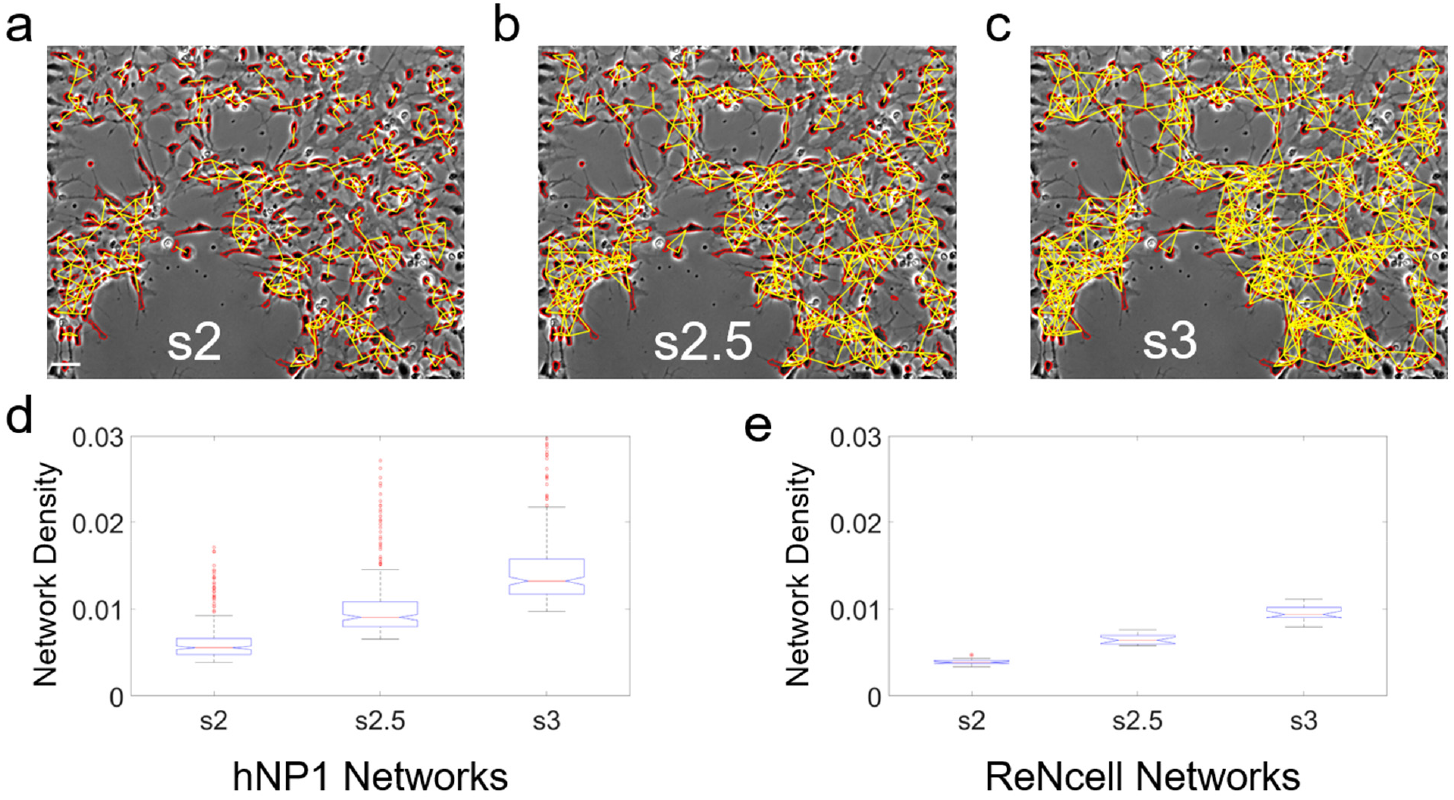
Scaling factor analysis. **(a-c)** hNP1 phase contrast image at day 3 of differentiation with graph representations using different scaling factors; scale bar = 50 μm. **(a)** scaling factor = 2; soma boundaries are outlined in red, and proximity edges are shown in yellow (scale bar = 50μm). **(b)** scaling factor = 2.5 **(c)** scaling factor = 3. **(d)** Boxplot of network density for all hNP1 networks. **(e)** Box plot of network density for all ReNcell networks, where nuclei were designated as nodes. Based on similarity in overall network density, scaling factor = 2 was used for phase contrast images (hNP1) and scaling factor = 3 was chosen for nucleus immunofluorescence images (ReNcell, NCRM-5 and CWI 4F2).

**Supplementary Figure 3.**
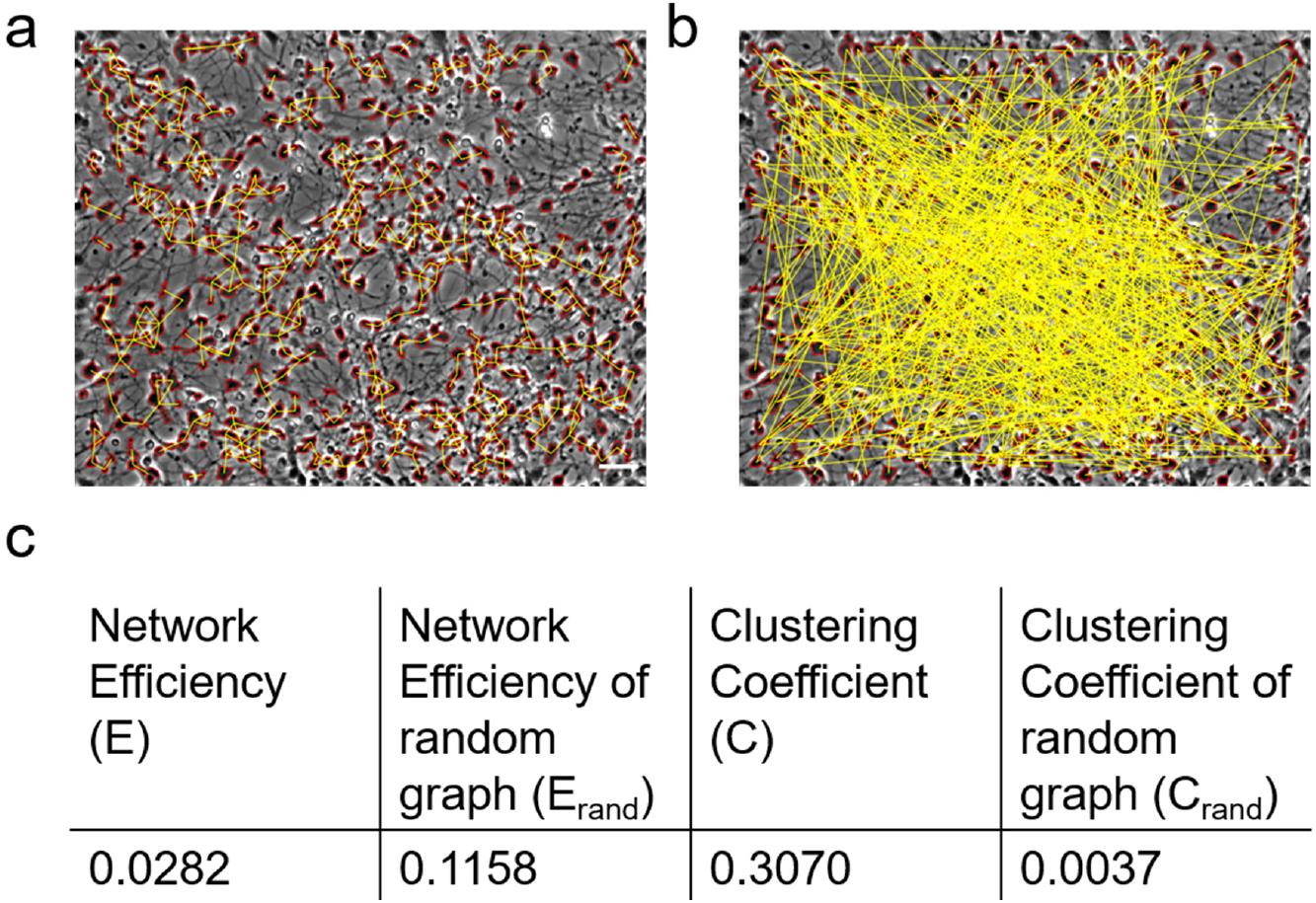
Random graph generation through degree-preserving rewiring. **(a)** Original graph representation of day 14 of differentiation image. Cell soma are outlined in red and edges are shown in yellow; scale bar = 50 μm. **(b)** Random graph generated by rewiring nodes in panel A with uniform probability. **(c)** Network parameters for graphs in (a) and (b).

**Supplementary Figure 4.**
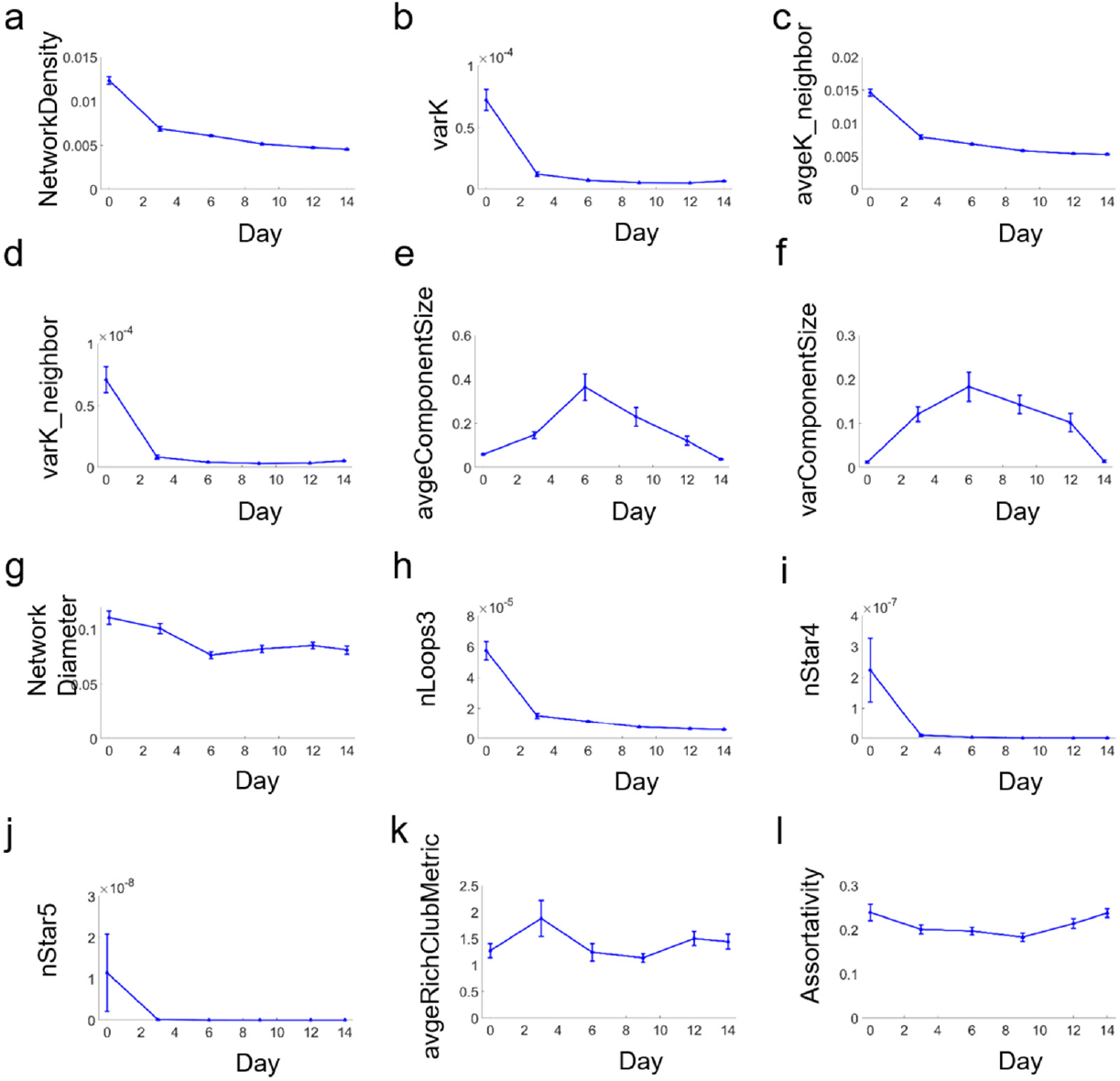
Trends in network metrics in hNP1 cultures not reported in main text. **(a-d)** All degree-related metrics decrease over 14 days of differentiation. **(a)** Network Density. **(b)** Variance in Degree. **(c)** Average Degree. **(d)** Variance in Neighbor Degree. **(e)** Average component size increases to a peak at day 6 and decreases as cells break up into smaller clusters. **(f)** Variance in Component Size is also highest at day 6. **(g)** Network Diameter. **(h-j)** Counts of loops and star motifs are highest at early stages, due to arrangement of neuroepithelial cells in rosette-like structures. **(h)** Triangular Loop Count. **(i)** 4-Star Motif Count. **(j)** 5-Star Motif Count. **(k)** Rich-Club Metric Average. **(l)** Assortativity.

**Supplementary Figure 5.**
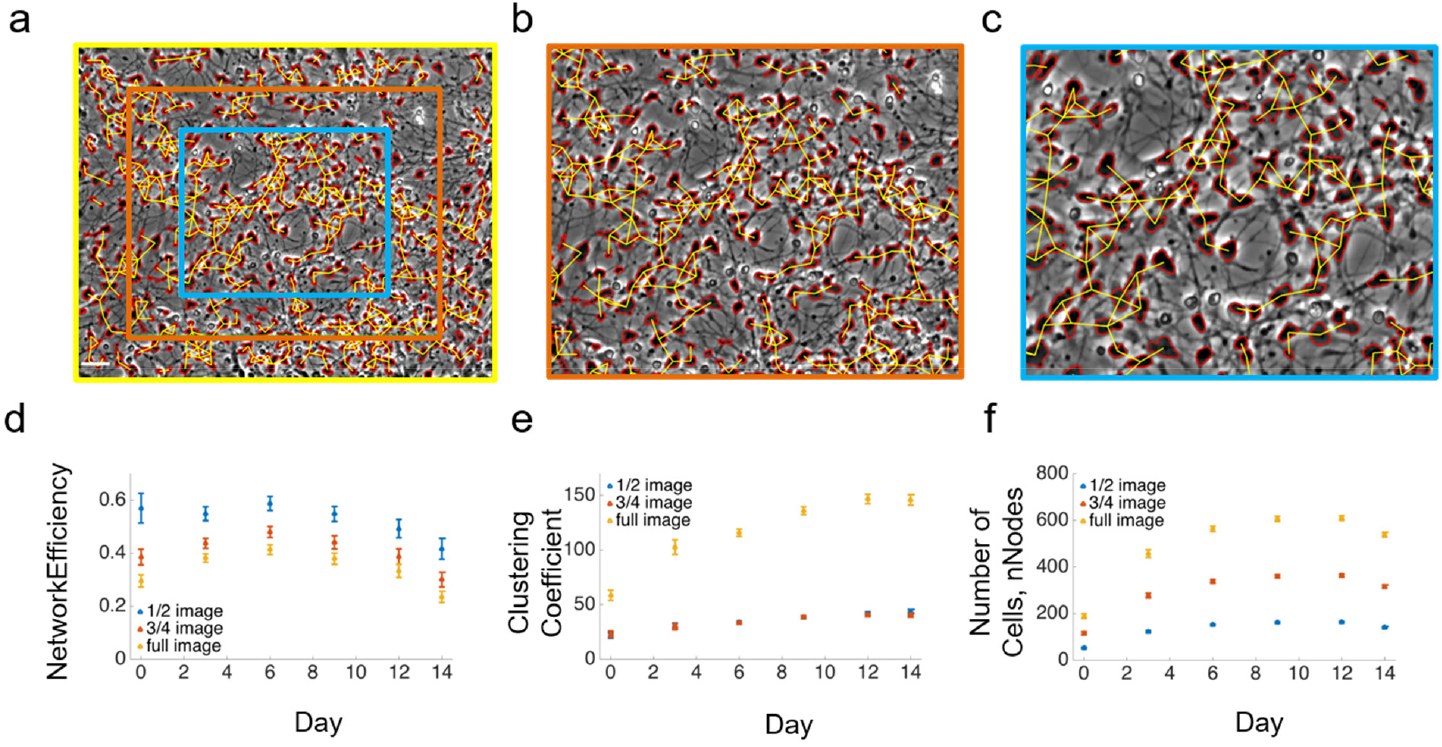
Scale invariance of network metrics in hNP1 network analysis. **(a-c)** Different fields of view chosen for network analysis. **(a)** Representative image at day 14 of differentiation; green box represents 75% of the field of view, white box represents 50% of the field of view (scale bar = 50μm). **(b)** Green inset from panel A. **(c)** White inset from (a). **(d-f)** Graph-based metrics computed for full image, 75% of the image and 50% of the image. **(d)** Network Efficiency across time. **(e)** Clustering coefficient. **(f)** Total number of cells in field of view. Values are reported as the mean across N = 30 networks ± SEM.

**Supplementary Figure 6.**
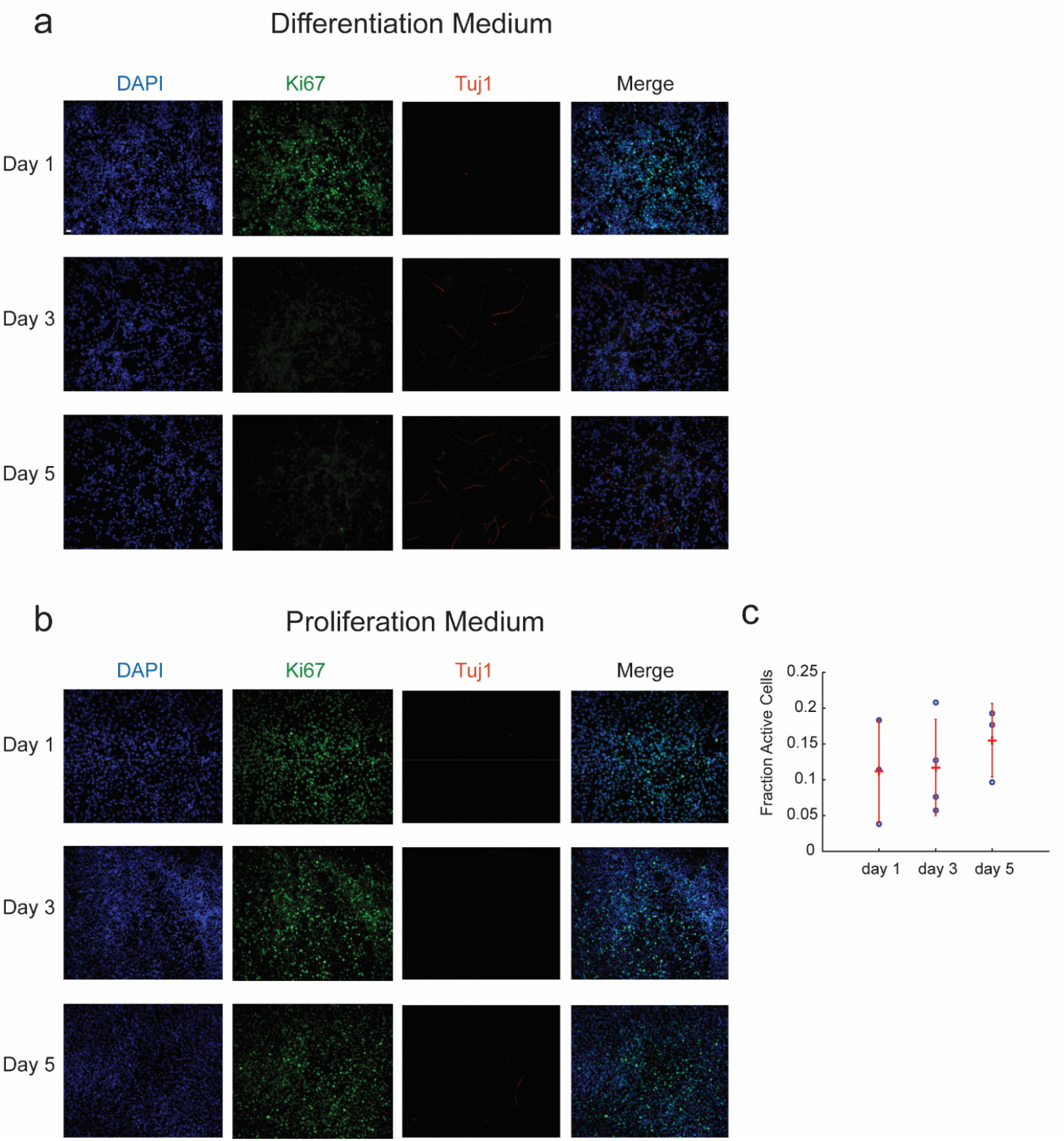
Immunostaining of ReNcell cultures and functional analysis of cultures with proliferation medium. **(a)** Immunostaining of cultures in differentiation medium for nuclei (DAPI), proliferating cells (Ki67) and new neurons (Tuj1); scale bar = 50μm. **(b)** Immunostaining of cultures in proliferation medium. **(c)** Fraction of active cells in cultures with proliferation medium (cells whose normalized fluorescence traces have three or more calcium transients).

**Supplementary Figure 7.**
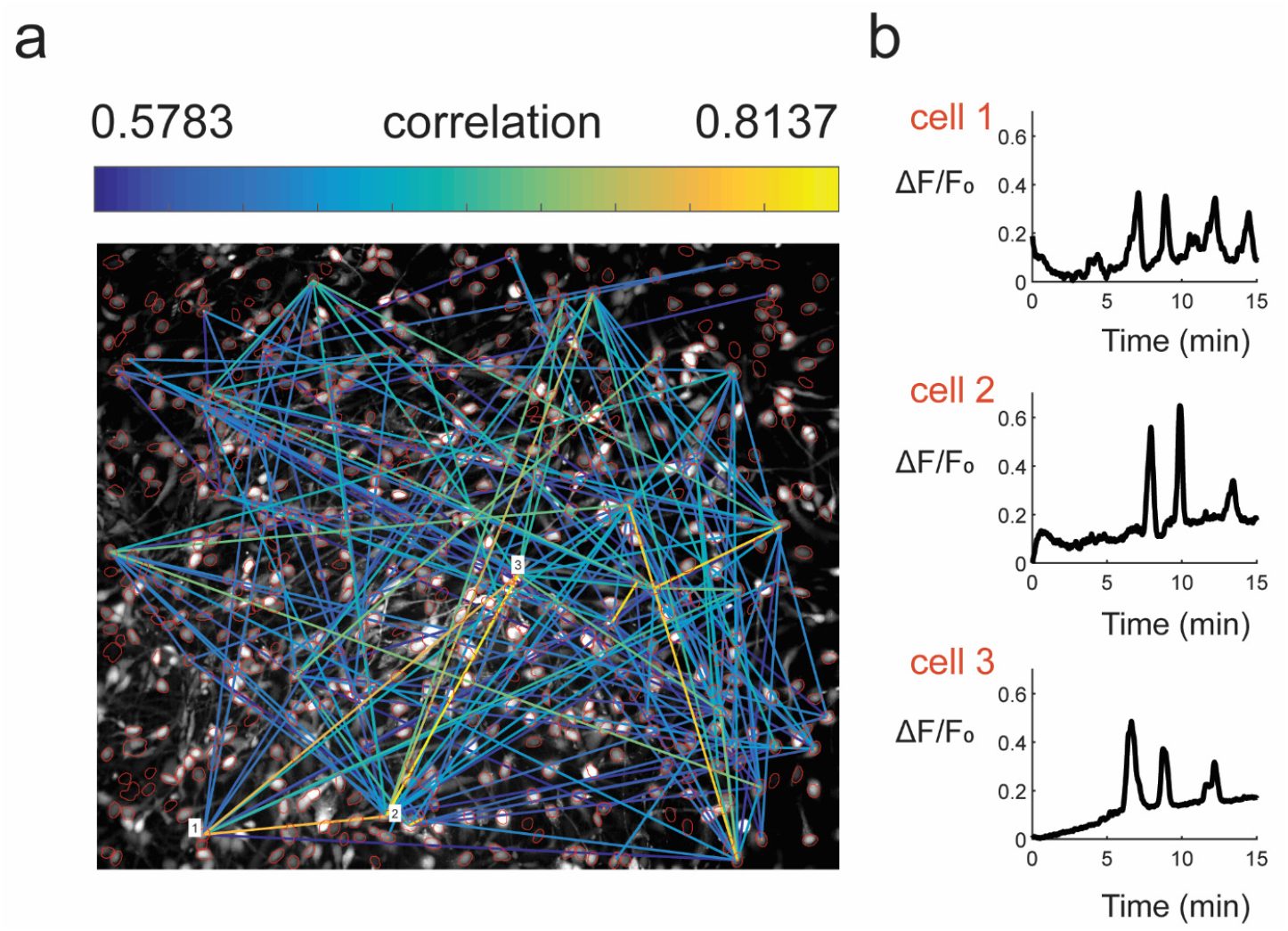
Cross-correlation analysis to infer functional connectivity from calcium imaging data. **(a)** Maximum intensity image from day 3 ReNcell culture loaded with Fluo-4 for calcium imaging. Inferred functional network is overlaid on the image, with correlation magnitude represented by edge color heatmap. ROIs obtained from corresponding nucleus image are shown in red. **(b)** Normalized calcium traces from 3 highly correlated cells marked in (a) are shown.

**Supplementary Figure 8.**
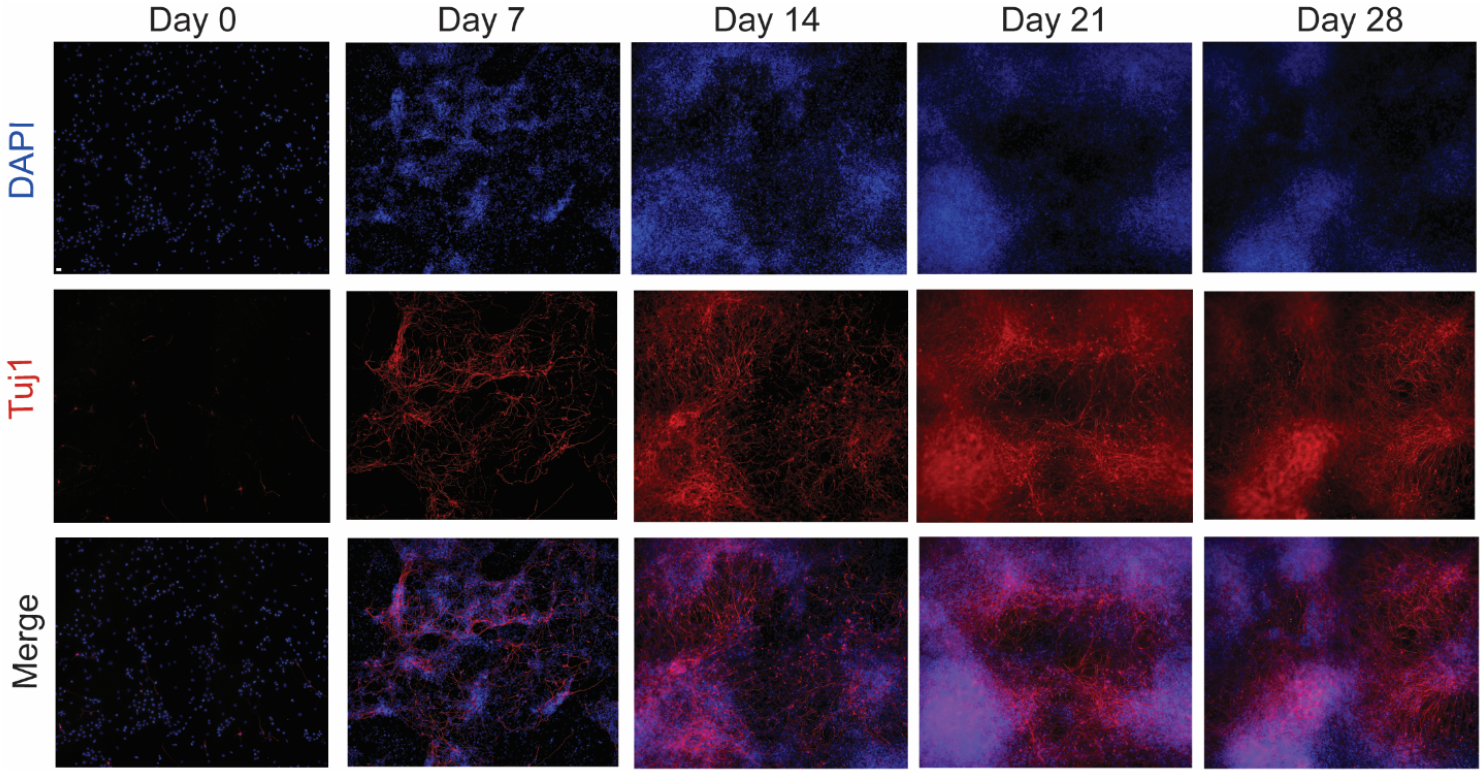
Long-term differentiation of NCRM-5 neural stem cells. NCRM-5 cultures at day 0, 7, 14, 21 and 28 stained for DAPI and β(III)-Tubulin (Tuj1). Cultures become more highly clustered at later stages of differentiation; scale bar - 50μm

**Supplementary Table 1.** List of parameters in image processing for hNP1 phase contrast images.

| Parameter Description | Parameter Value |
| --- | --- |
| Size of median filtering neighborhood | 3 pixels x 3 pixels |
| First threshold for removing small particles | 50 pixels |
| Second threshold for removing small particles (after separation of cell body objects) | 150 pixels |
| Radius of disk used for morphological opening | 4 pixels |
| Neighborhood connectivity used for connected component labeling | 8-connected neighborhood |

**Supplementary Table 2.** List of antibodies used for immunocytochemistry.

| Antibody | Host | Supplier | Catalog Number | Dilution |
| --- | --- | --- | --- | --- |
| Anti-Nestin | Mouse | Neuromics | MO15012 | 1:200 |
| Anti-MAP2 | Rabbit | Millipore | AB5622 | 1:500 |
| Anti-Tuj1 | Mouse | Millipore | MAB1637 | 1:2000 |
| Anti-Ki67 | Rabbit | Abcam | Ab15580 | 1.43μL/mL |
| Alexa Fluor 488 anti-mouse | Goat | Life Technologies | A-21121 | 1:1000 |
| Alexa Fluor 594 anti-rabbit | Goat | Life Technologies | A-11037 | 1:1000 |
| Alexa Fluor 594 anti-mouse | Goat | Life Technologies | A-11005 | 1:1000 |
| Alexa Fluor 647 anti-rabbit | Goat | Life Technologies | A-21244 | 1:1000 |

**Supplementary Table 3.** List of neural stem cell lines used in this study

| **Name** | **Source** | **Primary neuron type formed via differentiation** |
| --- | --- | --- |
| hNP1 | Derived from H9 human embryonic stem cells | - |
| ReNcell VM | Ventral mesencephalon of 10-week human fetal brain | Dopaminergic |
| NCRM-5 | Induced pluripotent stem cells derived from human male; NHCDR Cat# ND50031, RRID: CVCL_1E75 | Mixture of glutamatergic and GABAergic |
| CWI 4F2 | Induced pluripotent stem cells derived from female subject (6 months) with classical SLOS | Mixture of glutamatergic and GABAergic |

## Supplementary Videos

**Supplementary Video 1**. 8-day time-lapse movie of differentiating hNP1 human neural progenitor cells.

**Supplementary Video 2**. Same image sequence as in Supplementary Video 1, with cell bodies detected through image processing outlined in red and proximity edges shown in yellow.

**Supplementary Video 3**. Calcium imaging movie from day 1 ReNcell culture. Movie is sped up 100X. Original video was captured with a frame rate of 1Hz for a total duration of 15min.

**Supplementary Video 4**. Calcium imaging movie from day 3 ReNcell culture. Movie is sped up 100X. Original video was captured with a frame rate of 1Hz for a total duration of 15 min.

**Supplementary Video 5**. Calcium imaging movie from day 3 ReNcell culture showing calcium wave propagating through culture. Original video was captured with a frame rate of 1Hz for a total duration of 15 min.

**Supplementary Video 6**. Calcium imaging movie from day 5 ReNcell culture. Movie is sped up 100X. Original video was captured with a frame rate of 1Hz for a total duration of 15 min.

**Supplementary Video 7**. Four-day time lapse movie of differentiating ReNcell cultures. Video was captured at a frame rate of 1 frame/30min.

**Supplementary Video 8**. Calcium imaging movie from day 3 NCRM-5 culture. Movie is sped up 100X. Original video was captured with a frame rate of 1Hz for a total duration of 15 min.

**Supplementary Video 9**. Calcium imaging movie from day 3 NCRM-5 culture with immunostain image overlay showing Tuj1 (red) and Ki67 (blue) channels. Original video was captured with a frame rate of 1Hz for a total duration of 15 min.

**Supplementary Video 10**. Calcium imaging movie from day 7 NCRM-5 culture with immunostain image overlay showing Tuj1 (red) and Ki67 (blue) channels. Movie is sped up 100X. Original video was captured with a frame rate of 1Hz for a total duration of 15 min.

